# Enteroviral epitope mimicry enables NK cell-mediated targeting of ASPH in hepatocellular carcinoma

**DOI:** 10.64898/2026.04.07.717032

**Authors:** Man Hsin Hung, Qin Li, Limin Wang, Marshonna Forgues, Andrew S. Lee, Lisa M. Jenkins, Tapan K. Maity, Jesse Buffington, Jittiporn Chaisaingmongkol, Siritida Rabibhadana, Mathuros Ruchirawat, Mitchell Ho, Xin Wei Wang

## Abstract

Cancer development is shaped by host-microbe interactions, including viral infections. While several viruses are established oncogenic drivers, their potential protective roles in cancer remain unclear. Here we identify a dominant antibody response to CE1, a consensus epitope of enterovirus and rhinovirus, that is associated with reduced hepatocellular carcinoma (HCC) incidence and mortality. Anti-CE1 antibodies selectively recognize HCC cells and mediate anti-tumor activity through NK cell-mediated antibody-dependent cellular cytotoxicity (ADCC). Mechanistically, anti-CE1 antibodies cross-react with aspartate β-hydroxylase (ASPH), with CE1-ASPH sequence homology underpinning tumor recognition and cytotoxicity. Clinically, ASPH is aberrantly upregulated in HCC and correlates with inferred NK cell-associated ADCC activity and improved survival in CE1-seropositive patients. Collectively, these findings reveal a mechanism by which antiviral humoral immunity confers cancer protection through molecular mimicry and highlight anti-CE1 immunity as a potential therapeutic strategy in HCC.

## Introduction

Viruses exert complex roles in carcinogenesis and host immunity^1^. While oncogenic viruses, such as Epstein-Barr virus, can drive cellular transformation and promote tumorigenesis, emergent evidence also suggests viral infections may confer cancer-protective effects ^2–4^. For example, a positive serostatus for cytomegalovirus has been associated with enhanced T cell activation and improved clinical outcomes in melanoma patients receiving anti-PD1 checkpoint blockade^5^. In addition, spontaneous regression of leukemia and various solid tumors following acute viral infections has been reported^6,7^. Although the mechanism underlying virus-mediated anti-tumor effects remains unclear, one proposed model is that anti-viral immune responses cross-react with tumor antigens, thereby eliciting anti-tumor immunity. Supporting this concept, Rogone et al. reported that 60.6% of common tumor-associated antigens (TAAs) share sequence homology with viral peptides and demonstrated T cell cross-reactivity between TAA-viral-antigen pairs^8^. Similarly, Chiou et al. identified a tumor-associated epitope derived from TMEM161A in non-small cell lung cancer that cross-reacts with Epstein-Barr virus epitopes^9^. Moreover, GT103, an anti-complement-factor-H antibody derived from patients with early-stage non-small cell lung cancer, exhibits cross-reactivity with viral antigens from human metapneumovirus-1, human endogenous retrovirus, and measle virus^10^. Taken together, these findings highlight beneficial interactions between viral immunity and anti-tumor responses and support the potential of leveraging viral antigens to design novel cancer immunotherapies.

We explored beneficial interactions between viruses and cancer in hepatocellular carcinoma (HCC), the most common form of primary liver cancer and a leading cause of cancer-related mortality, responsible for an estimated 800,000 deaths annually worldwide^11^. Despite significant advances in immunotherapy and other targeted therapies, most patients with advanced HCC continue to experience therapeutic resistance and tumor progression^12,13^. New strategies are therefore needed to enhance immune-tumor engagement and improve anti-HCC immunity^14^.

Recently, we utilized pan-viral serological antibody profiling and identified that serological reactivity to rhinovirus (RV) and enterovirus (EV) was associated with improved clinical outcomes in patients with HCC^15^. EVs and RVs are ubiquitous, typically mild human pathogens. EVs comprise a large and diverse group of over 250 RNA virus types within the Picornaviridae family that infect multiple organ systems, whereas RVs, a respiratory-restricted subset with about 160 types across three species, primarily cause upper respiratory tract infections^16,17^. We demonstrated experimentally that a shared RV/EV epitope, CE1, can elicit T cell activation and promote CD8+ T cell-mediated killing of HCC cells^15^. However, it remained unclear whether the protective effects associated with EV/RV immunity are primarily mediated by the CE1 epitope and whether humoral immune responses targeting CE1 can also exert anti-HCC activity. Here, we addressed these questions by integrating structural analysis and seroreactivity profiling of EV/RV in HCC patients and healthy individuals. In addition, utilizing a phage-display human antibody scFv library, we identified anti-CE1 specific monoclonal antibodies and performed *in vitro* and *in vivo* studies to evaluate their anti-tumor activity and identify their molecular mimic targets in HCC. Our findings identify CE1 as a dominant epitope in EV/RV humoral immunity and uncover cross reactivity with aspartate β-hydroxylase (ASPH) as a mechanistic link between antiviral immunity and hepatocarcinogenesis. Enhancing anti-CE1 immunity may therefore offer a strategy for HCC therapy and prevention.

## Result

### CE1 defines protective EV/RV humoral immunity in HCC

EV and RV encode a single large polyprotein that is processed into structural capsid proteins (VP1, VP2, VP3, and VP4) and non-structural proteins with protease and RNA polymerase functions^18^. We found that CE1 is a consensus sequence located within the VP1 region (Figure 1a). To distinguish the contributions of CE1-specific and non-CE1 serological responses in HCC, we analyzed pan-viral serological antibody repertoire data from two independent cohorts^15,19^ (TIGER-LC, n=1,917; NCI-UMD, n=920). We identified serological responses to 2,011 viral peptides spanning 29 EV/RV strains with well-annotated structural information in UniProt^20^. We categorized the 2,011 viral peptides into three groups: 1) CE1-VP1, comprising 59 VP1 peptides overlap with the CE1 region; 2) non-CE1 VP1, comprising 281 peptides spanning the VP1 protein but excluding the CE1 region; and 3) non-VP1 EV/RV, comprising 1,671 peptides derived from EV/RV proteins other than VP1. We used normalized peptide counts derived from phage-immunoprecipitation-sequencing to calculate the epitope binding signal (EBS) as a measurement of serological reactivity for viral peptides. As shown in Figure 1a, seroreactivity to CE1-VP1 peptide was significantly higher than that of non-CE1 VP1 and non-VP1 EV/RV in both the TIGER-LC and NCI-UMD cohorts. Higher CE1-VP1 EBS was significantly associated with a reduced risk of HCC diagnosis (odds ratio= 0.9, *p*=0.015), whereas EBS for other EV/RV regions, as well as for Influenza and coronavirus peptides, showed no such trend (Extended Data Table 1). To further characterize EV/RV immune memory, we quantified the breadth of antibody response across epitopes of each EV/RV subcomponent using an antibody repertoire breadth (ARB) metric, calculated based on the normalized Shannon diversity index ^21^. CE1-VP1 exhibited a distinct distributional profile compared to non-CE1 VP1 and non-VP1 EV/RV (Figure 1b). Specifically, CE-VP1 ARB showed a consistent rightward shift, with most individuals exhibiting values >0.5, whereas non-CE1 VP1 and non-VP1 EV/RV ARB distributions were left-skewed, with values concentrated near zero. These results indicate that the antibody responses to CE1-VP1 are broadly distributed across different EV/RV strains, whereas responses to other EV/RV peptides are more restricted. Notably, CE1-VP1 ARB was significantly higher in healthy individuals than patients with HCC or chronic liver disease in both TIGER-LC (Chi-square *p*<0.00001) and NCI-UMD (Chi-square *p*=0.0121) cohorts (Figure 1c). Moreover, an ARB >0.5 for CE1-VP1 was significantly associated with improved overall survival in patients with HCC across both cohorts, independent of surgical eligibility (Figure 1d), while no survival association was observed for other EV/RV peptide groups (Extended Data Figure 1). Together, these findings suggest that the protective effects of EV/RV immunity in HCC are largely concentrated within the CE1 epitope.

**Figure 1.**
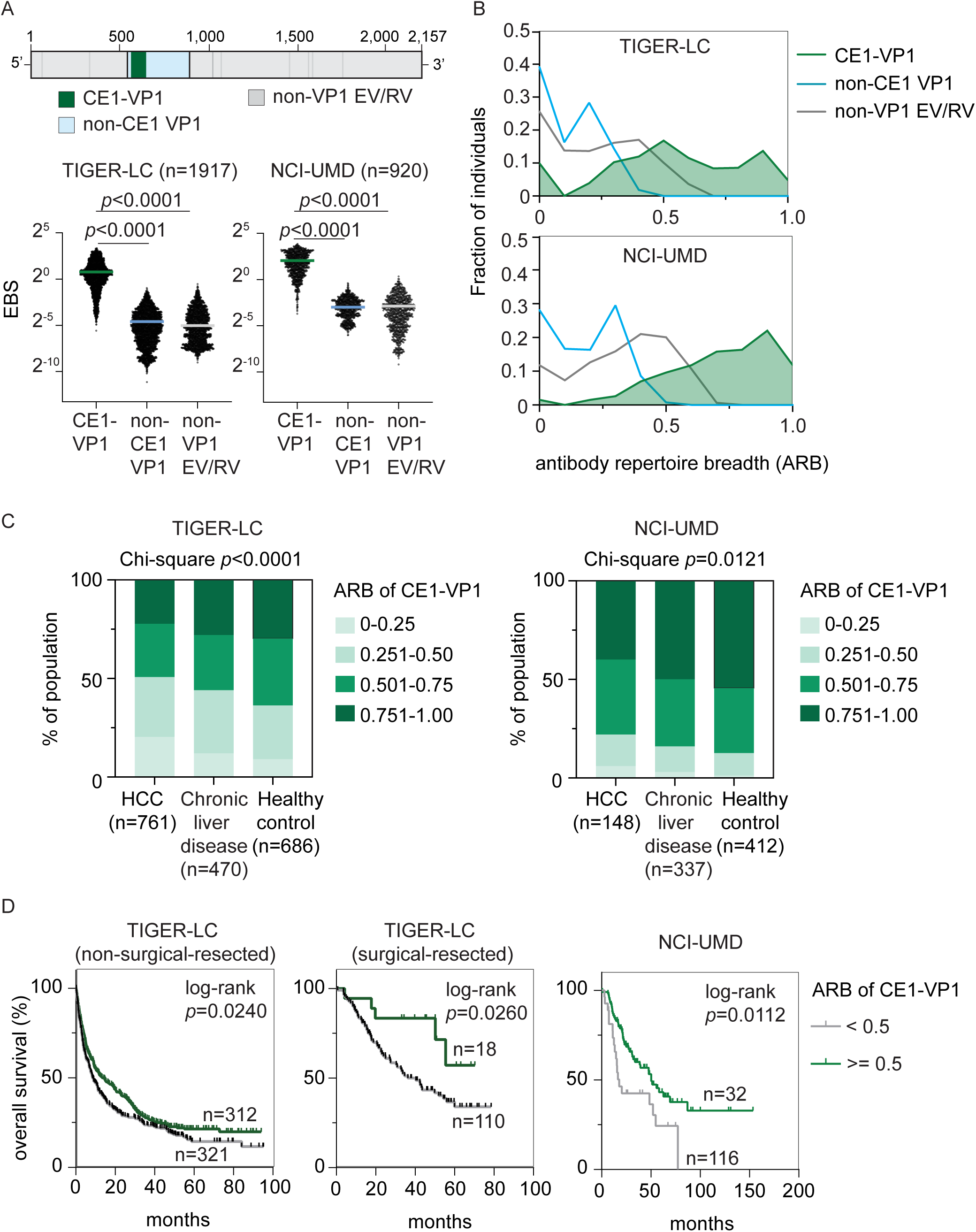
Distinct CE1-associated antibody response among EV/RV antibody repertoire is associated with clinical outcomes of HCC patients. (A) Average EBS level of viral peptides corresponding to each EV/RV subcomponent, namely CE1-VP1, non-CE1 VP1, and non-VP1 EV/RV (upper panel), in all individuals in the TIGER (n=1,917) and UMD (n=920) cohort. Bar, mean. Statistical significance was determined using a two-sided independent t-test. (B) Diversity of seroactive peptide among each EV/RV subcomponents was measured using antibody repertoire breadth (ARB) and the distribution of ARB of all individuals in TIGER and UMD cohort were summarized by histogram. (C) The distribution of ARB associated with CE1-VP1 in healthy volunteers, patients with chronic liver disease, and patients with HCC in TIGER and UMD cohort. ARB of CE1-VP1 were categorized into four groups as indicated and statistical significance was determined using Chi-square test. (D) Overall survival of HCC patients in TIGER cohort and UMD cohort stratified by ARB of CE1-VP1.

### Anti-CE1 antibodies recognize HCC cells

To determine whether CE1-VP1-specific antibodies exert anti-HCC activity, we screened a single-chain variable fragment (scFv) phage display library^22^ using a synthetic CE1 peptide conjugated with bovine serum albumin as the input antigen (Figure 2a). After three rounds of phage panning, enrichment of CE1-specific binders was observed (Extended Data Figure 2a). Colonies from the third round were isolated as single clones for subsequent binding specificity tests and sequence analysis. Five unique scFv clones were selected for recombinant expression, of which two (B9 and B10) were successfully expressed and exhibited specific binding to CE1 (Extended Data Figure 2b-c). The variable heavy and light chain domains of B9 and B10 were subsequently cloned into a rabbit immunoglobulin G gamma/kappa backbone to generate human-rabbit chimeric monoclonal antibodies, termed rB9 and rB10 (Figure 2a). Enzyme-linked immunosorbent assay (ELISA) confirmed that both antibodies bound the synthetic CE1 peptide in a dose-dependent manner (Figure 2b).

**Figure 2.**
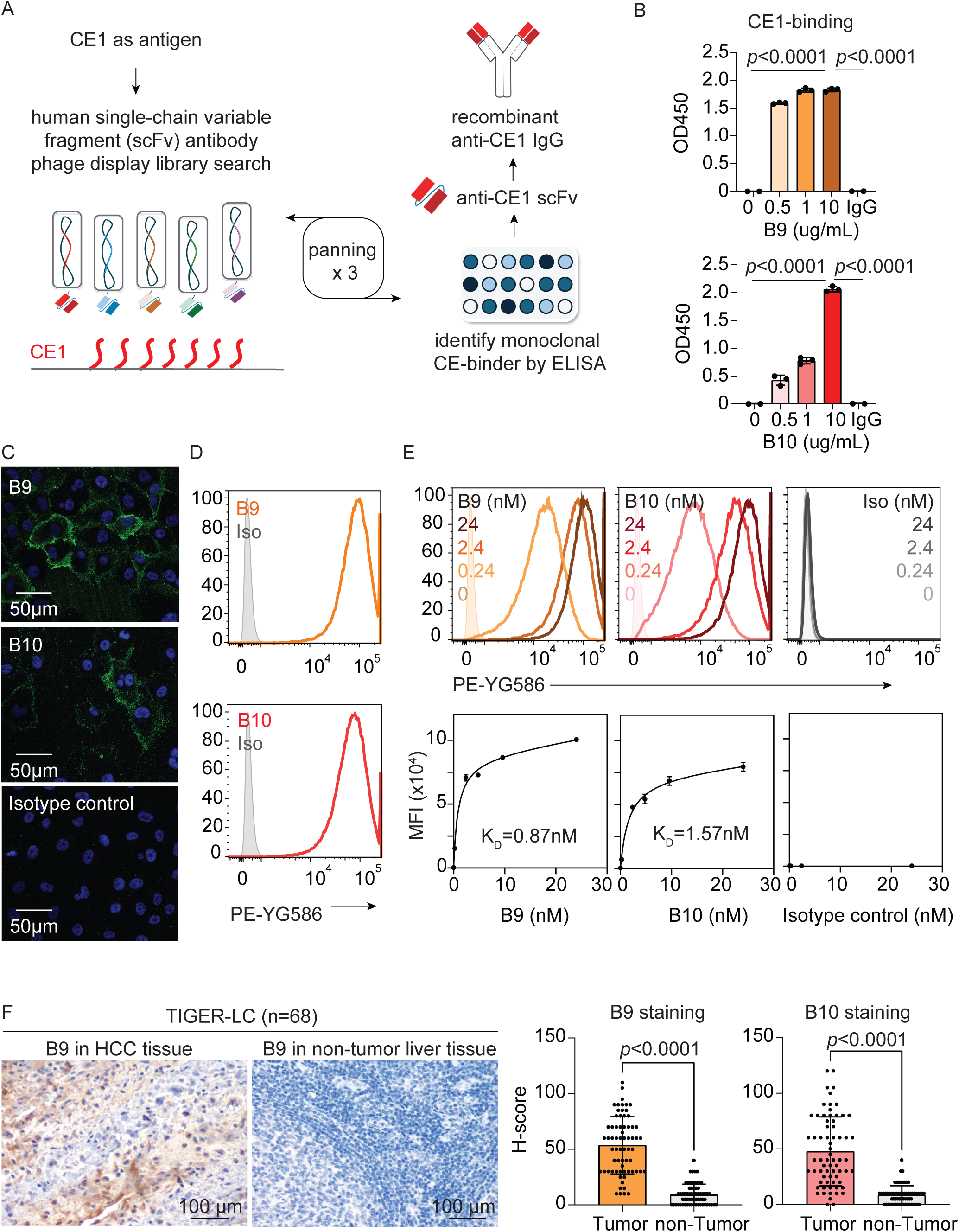
The generation and biological activities of synthetic monoclonal anti-CE1 antibodies, B9 and B10, in HCC. (A) Schematic summary from the identification of CE1-reacting single-chain variable fragments to the generation of synthetic monoclonal anti-CE1 antibody. (B) *Ex vivo* CE1 binding of B9 and B10. Individual data points are plotted over mean bar with error bars representing ± standard deviation (SD). Statistical significance was determined using a two-sided independent t-test. (C) Representative images of immunofluorescence staining of B9, B10 and control antibody in HLE cells. (D) Surface binding of B9, B10 and control antibody (2 μg/ml) to SNU475 cell. (E) Dose-dependent binding of B9, B10 and control antibody to SNU475 cell. (F) The binding of B9 and B10 to clinical HCC tumor and adjacent normal liver tissues were assessed using immunohistochemistry staining on samples obtained from 68 HCC patients in TIGER cohort. Average intensity of B9 and B10 in tumor and non-tumor tissues were shown at the right panel with representative images of B9 shown at the left panel. Individual data points are plotted over mean bar with error bars representing ± SD. Statistical significance was determined using a two-sided independent t-test.

Given the potential for viruses to influence host immunity via molecular mimicry^23^, we hypothesized that CE1 shares sequence homology with proteins expressed in HCC cells, thereby enabling cross-reactive binding. Immunofluorescence staining across five HCC cell lines (HLE, Huh7, SNU387, SNU398 and SNU475) revealed strong surface binding of rB9 and rB10, while no signal was observed with control antibodies (Figure 2c and Extended Data Figure 3a). We next extended this analysis to a panel of 18 HCC cell lines using flow cytometry. Consistent with immunofluorescence results, rB9 and rB10 exhibited significantly stronger binding than control antibodies across all HCC cells (Figure 2d and Extended Data Figure 3b). Dose-binding analysis in SNU475 cells yielded a K_D_ of 0.87 nM for rB9 and 1.57 nM for rB10, indicating high affinity binding (Figure 2e). Lastly, immunohistochemical analysis of tumor and non-tumor tissues from 68 HCC patients in the TIGER-LC cohort demonstrated significantly higher staining of rB9 and rB10 in tumor tissues (Figure 2f and Extended Data Figure 3c). Together, these results demonstrate that recombinant anti-CE1 antibodies (B9 and B10) exhibit high-affinity, HCC-specific surface recognition.

### Anti-CE1 antibodies mediate antibody-dependent cellular cytotoxicity in HCC

To elucidate the therapeutic potential of anti-CE1 antibodies in HCC, we generated fully humanized versions of B9 and B10 by cloning their variable domains into a human IgG1 backbone and expressing them in Human Embryonic Kidney 293 (HEK293) cells. Specific binding of humanized B9 (hB9) and B10 (hB10) to CE1 was confirmed by ELISA (Extended Data Figure 4). Antibody-dependent cellular cytotoxicity (ADCC) mediated by natural killer (NK) cells is a key mechanism underlying antibody-based anti-tumor activity ^24^. To examine whether anti-CE1 antibodies can induce ADCC against HCC, we performed co-culture assays in which HCC cells were incubated with antibodies and NK cells, and tumor cell growth was monitored using a label-free, impedance-based system ^15^. As shown in Figure 3a, hB9 and hB10 in the presence of NK cells significantly inhibited the growth of multiple HCC cell lines (HLE, Huh7, SNU387, and SNU475) compared with control antibody. This effect was dose-dependent and was not observed with the control antibody (Figure 3b). To further evaluate *in vivo* activity, we utilized nude mice, which retain high NK cell activity^25^. Mice bearing subcutaneous Huh7 xenografts were treated with hB9, hB10, or control human IgG antibody (10 μg/kg) via tail-vein injection three times per week. Compared with mice receiving control antibody, the Huh7 tumor growth in both hB9- and hB10-treated mice was significantly suppressed (Figure 3c-d), without affecting body weight (Figure 3e) or liver function (Extended Data Figure 5). Collectively, our data showed that humanized anti-CE1 antibodies elicit potent NK cell-mediated ADCC against HCC cells *in vitro* and *in vivo*.

**Figure 3.**
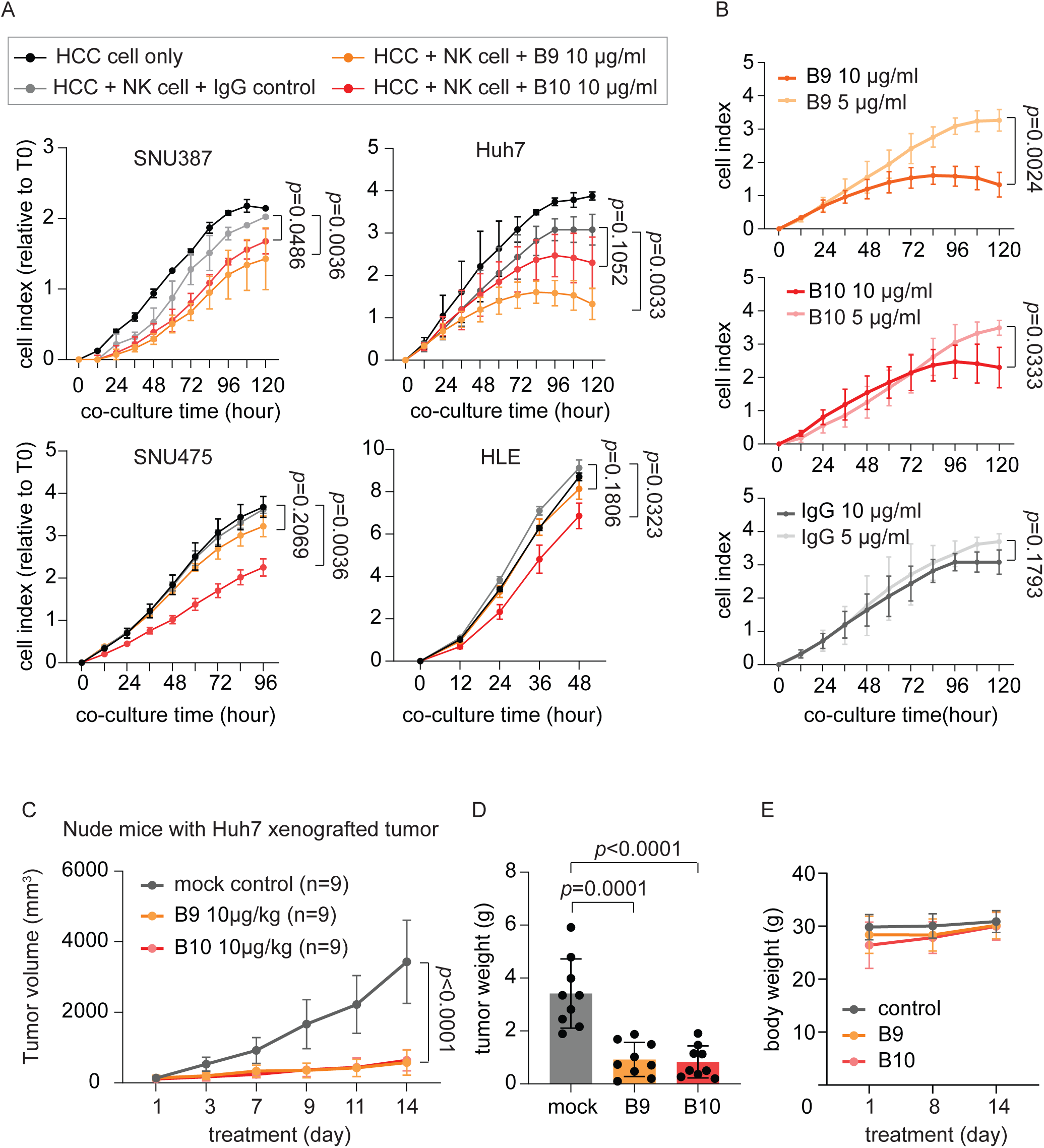
B9 and B10 promote NK cell-mediated ADCC against HCC *in vitro* and *in vivo*. (A) Growth curves of SNU387, Huh7, SNU475 and HLE cells after incubating with NK cells plus B9, B10, or control antibody. (n=3) HCC cells were seeded at the density of 4,000 cell per well on a 16-well E-plate, followed by adding 4,000 NK cells plus antibody (10 μg/ml) per well on the other day. Confluence of HCC cells was continuously monitored using xCELigence and average growth signal at indicated time were shown as dots with error bars representing ± SD. Statistical significance was determined using a two-way ANOVA test. (B) Dose-dependent ADCC effects of B9, B10 and control antibody in Huh7. (n=3) (C) The *in vivo* anti-HCC effects of B9, B10 and control antibody. Huh7 xenografted tumors were established on Nude mice and antibody treatments (10 μg/kg, three times a week) were given via tail vein injection when xenografted tumors reached around 100 mm^3^. Tumor size at indicated time were summarized as dot with error bars representing ± SD. Statistical significance was determined using a two-way ANOVA test. (D) Average weight of xenografted Huh7 tumors after B9, B10 and control antibody treatment. Individual data points are plotted over mean bar with error bars representing ± SD. Statistical significance was determined using a two-sided independent t-test. (E) Average body weight of mice under B9, B10 and control antibody treatment. Body weight at indicated time were summarized as dot with error bars representing ± SD.

### Identification and validation of ASPH as a target of anti-CE1 antibodies in HCC

The above results suggested that B9 and B10 recognize endogenous cell surface proteins in HCC. To identify their targets, we performed immunoprecipitation followed by mass spectrometry using rB9, rB10 and control antibodies on cell lysates from Huh7 and HLE cells. Antibody-associated complexes were analyzed by mass spectrometry using data-independent acquisition, enabling unbiased identification of binding partners^26^ (Figure 4a). Comparative analysis identified 1454 candidate proteins that were significantly enriched by rB9 or rB10 (>100-fold over control) in both cell lines. Cross-referencing these candidates with a curated human surface proteome database ^27^ yielded 47 surface proteins. Given that B9/B10-binding is likely driven by sequence similarity to CE1, we evaluated sequence homology and associated expected value of alignment^28^, which identified ASPH as the top candidate (Figure 4b-c).

**Figure 4.**
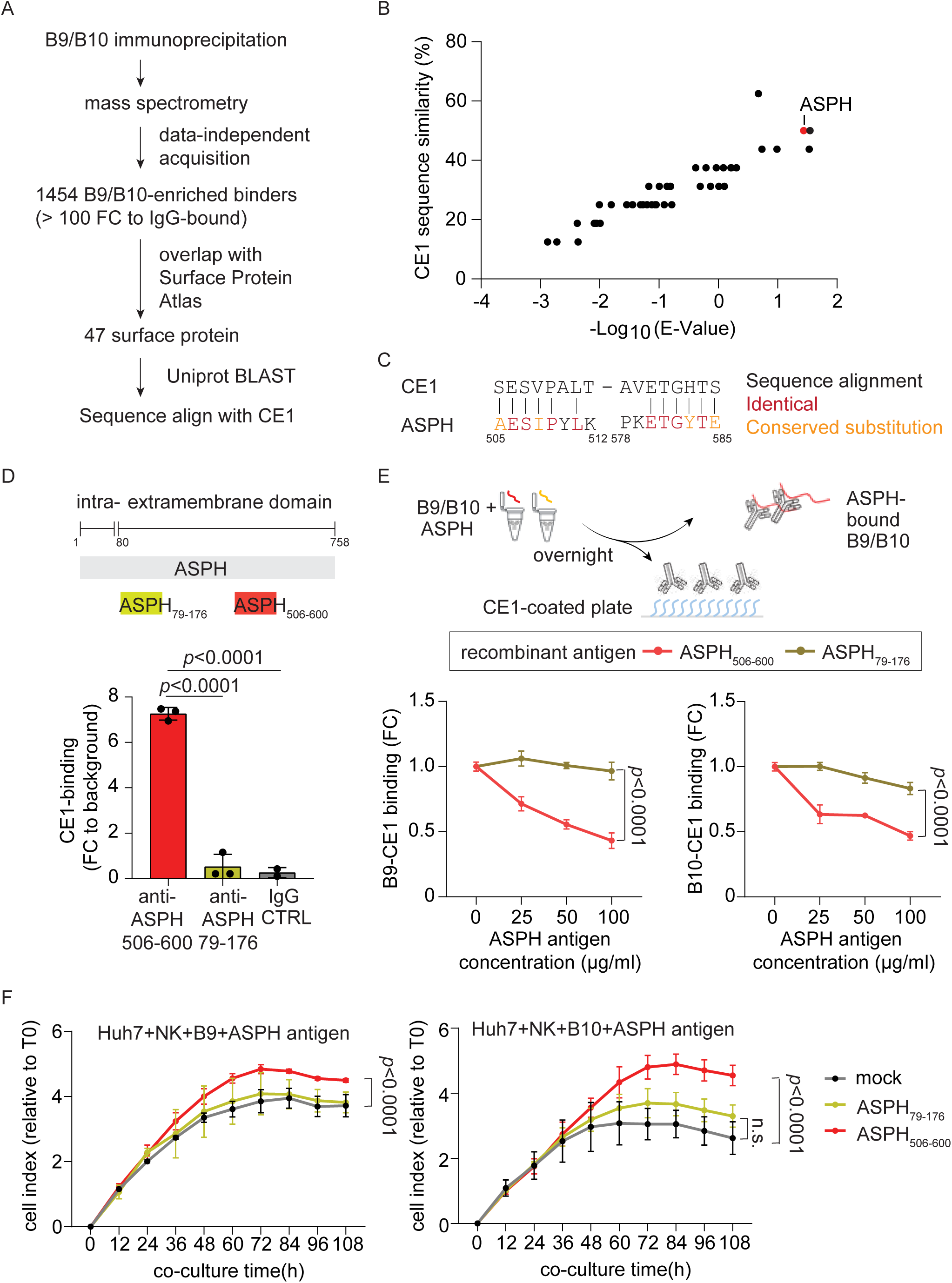
B9 and B10 cross react with human ASPH, mediating the anti-HCC effects. (A) Schematic summary of the workflow combining immunoprecipitation-mass spectrometry study and bioinformatic strategies to identify human surface protein targeted by B9 and B10 in HCC. (B) Sequence alignment analysis compares the sequence similarity of the 47 human surface proteins bound by B9/B10 with CE1. (C) Sequence alignment of CE1 and ASPH. Amino acid pair with a BLOSUM62 score larger than 0 is considered as conserved substitution. (D) Ex vivo CE1-recognition by anti-ASPH antibody. Upper panel illustrated the locations of known antibody-recognition-sites of ASPH, namely ASPH_79-176_ and ASPH_506-600_. Antibodies targeting ASPH_79-176_ and ASPH_506-600_ were incubated with synthetic CE1 peptide for two hours and the binding between CE1 and anti-ASPH antibodies were assessed using ELISA. An isotype control was included for the test. (n=3). Individual data points are plotted over mean bar with error bars representing ± SD. Statistical significance was determined using a two-sided independent t-test. (E) Competition of recombinant ASPH antigens with CE1 peptide to CE1-B9/B10 binding. B9 and B10 (2 μg/ml) were pre-incubated overnight with recombinant ASPH_79-176_ and ASPH_506-600_ at 25, 50, and 100 μg/ml (approximately 2.5-, 5-, and 10-fold molar excess relative to B9/B10 antibody). Subsequently, CE1-binding by B9 and B10 were assessed using ELISA. Average binding signal at indicated conditions were summarized as dot with error bars representing ± SD. Statistical significance was determined using a two-way ANOVA test. (F) Effects of recombinant ASPH antigens in B9/B10-mediated anti-HCC activities. First, Huh7 cells were seeded at the density of 4,000 cell per well on a 16-well E-plate, and B9/B10 (5 μg/ml) were incubated with recombinant ASPH antigens ASPH_79-176_ and ASPH_506-600_ at 125 μg/ml. After overnight incubation, B9/B10-ASPH mixture plus 4,000 NK cells per well were added to Huh7 cells, and the growth of Huh7 cells was continuously monitored using xCELigence. Average growth signal at indicated time were shown as dots with error bars representing ± SD. Statistical significance was determined using a two-way ANOVA test.

ASPH contains two reported antigen recognition sites (amino acids 79-176 and 506-600)^29,30^, with the latter overlapping the CE1-mimicking region (Figure 4c-d). To validate ASPH as a target, we tested commercially available antibodies against these regions. Anti-ASPH_506-600_ showed significantly stronger binding to the CE1 peptide than anti-ASPH_79-176_ and control antibody, consistent with the sequence alignment results (Figure 4d). To further confirm CE1-ASPH mimicry, we generated recombinant ASPH fragments (amino acids 79-176 and 506-600, each fused to an albumin-tag; termed as rASPH_79-176_ or rASPH_506-600_, respectively) and verified their antigenicity by ELISA (Extended Data Figure 6). Pre-incubation of rB9 and rB10 with different doses of rASPH_506-600,_ but not rASPH_79-176_, dose-dependently inhibited their binding to CE1 (Figure 4e), indicating that this region mediates the interaction. Lastly, to assess functional relevance, recombinant rASPH_79-176_ and rASPH_506-600_ were pre-incubated with hB9 or hB10 overnight and tested in NK cell-mediated ADCC assays using Huh7 cells. rASPH_506-600_, but not by rASPH_79-176_, significantly attenuated B9/B10-mediated ADCC (Figure 4f). Together, these results demonstrate that ASPH, specifically its 506-600 region, acts as a molecular mimic of CE1 and mediates both antigen recognition and anti-tumor activity of anti-CE1 antibodies in HCC.

### ASPH is aberrantly expressed in HCC and associates with NK cell activity and overall survival in CE1-seropositive patients

To investigate the role of ASPH in HCC, we analyzed four transcriptomic datasets associated with three independent clinical cohorts, including TCGA-LIHC (tumor, n=373; non-tumor, n=50), LCI (tumor, n=247; non-tumor, n=239), and TIGER-LC (set 1: expression array; tumor, n=62; non-tumor, n=59; set 2: RNA-seq; tumor, n=51; non-tumor, n=51)^31,32^. Across all datasets (733 tumors and 399 non-tumor tissues), ASPH expression was consistently elevated in HCC tumors compared with non-tumor liver tissues (Figure 5a). Single-cell transcriptomic analysis further confirmed that ASPH expression was largely restricted to malignant cells and was minimally detected in non-malignant cells (Figure 5b and Extended Data Figure 7).

**Figure 5.**
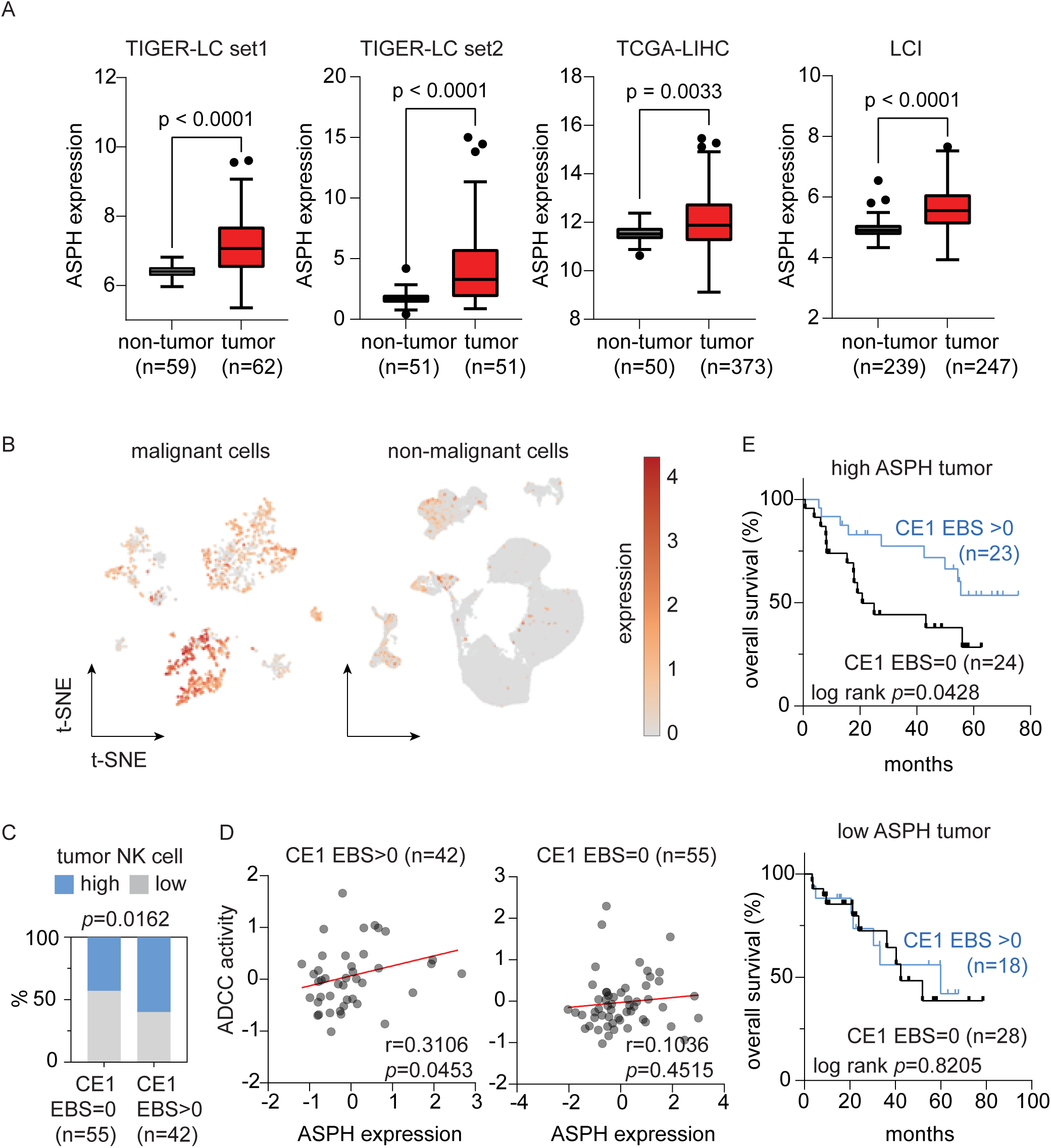
ASPH is upregulated in HCC and associated with NK cell activity and clinical outcomes in CE1 active patients. (A) Box plots summarize the expression of ASPH in clinical HCC tumor tissues and non-tumor liver tissues. The central rectangles span from the first quartile to the third quartile, with lines inside the rectangles indicating the median and whisker extending 1.5 times the interquartile range. Statistical significance was determined using a two-sided independent t-test. (B) t-SNE plots show the expression of ASPH in liver cancer cells (n=1,574) and non-malignant cells (n=77,670). (C) Bar plot summarize the percentage of high- and low-NK-cell-infiltrated tumors in patients with and without detectable CE1 seroreactivity. Level of tumor-infiltrating NK cell was estimated using xCell^33^, and EBS of the 40 most reactive CE1-related EV/RV viral peptides^15^ were used to determine CE1 seroreactivity. Statistical significance was determined using Chi-square. (D) The relationship between tumor ASPH expression and ADCC activity. Statistical significance was determined using Spearman correlation test. (E) Kaplan-Meier curve plots summarize the overall survival of HCC patients according to their CE1 seroreactivity and tumor ASPH expression in TIGER-LC cohort. Median ASPH expression in the HCC tumors and the EBS of the 40 CE1 peptides were used for stratification. Statistical significance was determined using log-rank test.

To determine the interplay between CE1 seroreactivity, ASPH expression and NK cell activity, we analyzed 97 HCC patients from the TIGER-LC cohort with matched tumor transcriptomic and pan-viral serology data. Tumor deconvolution analysis^33^ revealed significantly higher NK cell infiltration in patients’ tumors with detectable CE1 seroreactivity (p=0.0162, Figure 5c). Furthermore, ASPH expression was positively correlated with inferred ADCC activity^34^ in tumors from CE1-seropositive patients (p=0.0453), while no such association was observed in CE1-seronegative tumors (p=0.4515, Figure 5d). We next assessed whether interactions between ASPH expression and CE1 seroreactivity are associated with clinical outcomes of patients with HCC. Among patients with high ASPH expression, those with detectable CE1 seroreactivity exhibited improved overall survival compared with those without (p=0.0428). In contrast, CE1 seroreactivity was not associated with overall survival in patients with low ASPH expression (Figure 5e). Collectively, these results demonstrate that ASPH is aberrantly upregulated in HCC and functionally links with CE1-associated immunity to NK cell activity and clinical outcomes.

## Discussion

In this study, we interrogated the anti-HCC activity of EV/RV antibodies in two independent clinical cohorts comprising 909 HCC patients and 1098 healthy individuals. In our previous work, we identified a shared EV/RV epitope, CE1, that elicits CD8+ T cell activation and promotes T cell-mediated killing of HCC cells^15^. Building on this finding, we screened through 2,011 unique peptides spanning the full proteomes of 29 EV/RV species and identified an antibody response to CE1, a consensus epitope within the VP1 capsid protein, that exhibited a distinct profile associated with anti-HCC properties. The antibody response to CE1 was both stronger and more diverse than the responses to other regions of EV/RV, despite the presence of antigenic sites elsewhere in these proteins^35,36^. Consistent with this, both the magnitude and diversity of CE1-specific antibody responses across EV/RV species were associated with reduced risk of HCC diagnosis and lower HCC-related mortality, whereas such associations were not observed for antibodies targeting non-CE1 EV/RV proteins or other respiratory viruses.

The generation, preservation, and propagation of antibody repertoire are shaped by a balance of host benefit and metabolic costs^37,38^. Chardès et al. demonstrated that following repeated viral exposure, the immune system suppresses de novo antibody maturation when existing memory provides sufficient protection thereby minimizing metabolic plasticity cost^38^. The quantitative and qualitative dominance of CE1-specific responses therefore suggests selection pressure shaping the EV/RV antibody repertoire. Although the precise mechanisms underlying this selection remain unclear, we propose that the CE1-associated antibody clones may confer additional fitness benefits beyond antiviral protection, contributing to their immunodominance. Recent epidemiological studies have reported reduced risks of neurodegenerative diseases following influenza vaccination^39,40^ and herpes zoster vaccination^41,42^. Given the distinct biology of these viruses, such neuroprotective effects are unlikely to be solely attributable to antiviral activity. Together with our findings, these observations support the concept that certain virus-associated antibodies may confer benefits beyond infection control, thereby enhancing host fitness. They also highlight the potential of leveraging anti-viral antibodies as therapeutic agents in non-viral infectious diseases, including cancer.

In this study, we generated two monoclonal anti-CE1 antibodies and experimentally demonstrated their ability to recognize HCC cells and promote NK cell-mediated ADCC. The results provide direct evidence for cross-reactivity between CE1-directed humoral immunity and HCC, supporting our clinical observations. Importantly, we identified ASPH as a molecular mimic target linking anti-CE1 antibodies to tumor recognition and cytotoxicity. ASPH is a Type II transmembrane protein that catalyzes the hydroxylation of aspartyl and asparaginyl residuals within epidermal-growth-factor-like domains of multiple proteins, including Notch, and plays key roles in oncogenic signaling pathways regulating cell growth, motility and invasiveness^43 44^. ASPH also exerts catalytic independent oncogenic functions, such as activation of SRC signaling through interaction with ADAM12/ADAM15^45^. Clinical studies showed that ASPH is frequently upregulated in a wide range of malignancies, including HCC and pancreatic cancer, but is minimally expressed in normal adult tissues, and elevated ASPH expression is associated with tumor differentiation and adverse clinical outcomes^43,46^. Notably, ASPH is being actively explored as a potential therapeutic target. Small-molecule ASPH inhibitors were found to suppress tumor growth by inhibiting ASPH-regulated signaling pathways in preclinical models^44,45^. Meanwhile, ASPH has been developed as a cancer-associated antigen for monoclonal antibody, vaccine, and chimeric antigen receptor-T cell therapies ^47–49^. In our study, anti-CE1 antibodies showed preferential binding to HCC cells compared with adjacent non-malignant liver tissues and were capable of eliciting NK cell-mediating ADCC, consistent with the known expression pattern and therapeutic relevance of ASPH. An immunoprecipitation-mass spectrometry study confirmed the interaction between ASPH and anti-CE1 antibodies, and an in vitro competition assay delineated the ASPH epitope responsible for anti-CE1-mediated recognition and anti-HCC effects. Integrative study of tumor transcriptomes and viral serology further supported the interaction between CE1 seroreactivity and ASPH level expression, linking to NK cell activity and clinical outcomes. Collectively, our findings support ASPH as a molecular mimic target mediating the anti-HCC effects of EV/RV-induced humoral immunity. Given the high prevalence of EV/RV globally and aberrant activation of ASPH in multiple cancers, it will be of interest to investigate whether the EV/RV-associated protection and anti-CE1 antibody-based strategies extend to other malignancies. In addition, it remains unclear whether ASPH also mediates CE1-specific CD8+T cell responses, which warrants further investigation.

In this study, we demonstrated the cross-reactivity between EV/RV-induced humoral immunity and HCC using anti-CE1 antibodies. CE1 is a consensus sequence derived from 40 reactive VP1 peptides^15^, incorporating the most conserved residues across these sequences, although no individual native peptide is fully identical to CE1. This approach preserves the diversity of native VP1 proteins. For a therapeutic perspective, introducing antibody targeting CE1 rather than individual viral epitopes may increase antibody repertoire diversity, which in our study was associated with improved clinical outcomes in patients with HCC.

However, this study has several limitations. The use of recombinant CE1-based antibodies may not fully capture the complexity of naturally occurring antibody responses, as the phage-display screening was performed using a single linear CE1 peptide. Although the CE1-overlapping VP1 region is largely linear^50,51^, this approach may miss antibodies recognizing conformational or multi-region epitopes. In addition, variability in affinity among endogenous EV/RV antibodies toward ASPH and HCC cannot be excluded. Our analyses also focused on IgG-based antibodies, whereas natural responses may involve multiple isotypes. Finally, EV/RV-associated anti-tumor effects may extend beyond humoral immunity to include cellular and innate mechanisms. Consistent with this, our previous work demonstrated CE1-mediated CD8 T cell activation, and other studies have shown that SARS-CoV-2 virus triggered the expansion of CCR2+ nonclassical monocytes via releasing its viral RNA and attenuated tumor progression in metastatic murine tumor models associated with melanoma, lung, breast, and colon cancer^52^. Further studies are required to define the broader immune mechanisms underlying EV/RV-associated cancer protection.

## Method

### Clinical cohorts

For pan-viral serological antibody repertoire analysis, we identified 2,837 participants from two independent cohorts, TIGER-LC (n=1,917) and NCI-UMD (n=920), with available viral profiling and clinical data reported in previous studies^15,19,53^. The TIGER-LC cohort comprised HCC patients (n=761), patients with chronic liver diseases (n=470), and healthy individuals (n=686). The NCI-UMD cohort included HCC patients (n=148), patients with chronic liver diseases (n=337), and healthy individuals (n=412). For bulk tumor transcriptomic analysis, we interrogated 733 HCC samples and 399 tumor-free liver tissue samples from three independent published cohorts, specifically included 247 tumors and 239 associated non-tumor tissues from LCI cohort (GSE14520)^54^, 373 tumors and 50 associated non-tumor tissues from TCGA-LIHC cohort (TCGA Research Network: https://www.cancer.gov/ccg/research/genome-sequencing/tcga)^55^, and 113 tumors and 110 non-tumor tissues from TIGER cohort (dbGaP Study Accession phsphs001199.v2.p1)^31,56^. For single cell analysis, we utilized the publicly available data from scAtlasLC (https://scatlaslc.ccr.cancer.gov/#/), comprising 222,862 single-cell transcriptomic profiles associated with 101 liver cancer patients from three independent cohorts^57–59^. Additionally, 68 HCC patients with available tumors and non-tumor tissues from TIGER-LC were identified for an immunohistochemistry (IHC) study.

### Enteroviral (EV) and rhinoviral (RV) antibody repertoire analysis

EV/RV antibody responses were obtained from previously published pan-viral serological profiling datasets generated using phage-immunoprecipitation-sequencing^15,19,53^. Briefly, 2 mg of total immunoglobulins per serum sample was used for immunoprecipitation with a bacteriophage library displaying 93,904 viral epitopes from 206 human pathogenic viral species. Signals from precipitated phages were measured using the Illumina NextSeq 500 sequencing platform at a depth of 200 million reads per sequence lane. The epitope binding signal (EBS), defined as the enrichment Z-score of read counts, was used as a relative quantification of antibody reactivity. To determine how epitope location influences antibody reactivity, we analyzed antibody responses of 2,011 EV/RV viral epitopes derived from 29 EV/RV polyproteins (UniPort IDs: P03300, P03303, P03313, P04936, P06210, P07210, P12916, P21404, P23008, P23069, P29813, P36290, Q03053, Q66474, Q66575, Q66577, Q66790, Q82081, Q82122, Q9WN78, Q9YLG5, Q9YLJ1, P08291, Q66282, Q86887, P08292, Q9QL88, P03301, and P03302). Structure annotations of the 29 EV/RV polyproteins, including the start and end coordinates for VP1-4 protein and other non-structure proteins, were obtained from the UniProtKB database^20^ using the Chain identifier. Based on the structure annotation and overlap with the CE1 region, the 2,011 EV/RV epitopes were categorized into three groups: 1) CE1-VP1 (n= 59), comprising VP1 epitopes overlapping with CE1; 2) non-CE1 VP1 (n= 281), comprising VP1 epitopes excluding CE1; and 3) non-VP1 EV/RV (n= 1,671), comprising epitopes derived from regions outside VP1. Additionally, EBS of 2178 viral epitopes associated with coronaviruses and EBS of 4635 viral epitopes associated with Influenza viruses were obtained. Logistic regression was performed using the glm() function in R (version 4.5.1) to model the association between antibody reactivities and risk of HCC diagnosis across EV/RV, coronavirus and influenza. To compare the breadth of antibody responses across viral EV/RV epitopes, we calculated an antibody repertoire breadth metric (ARB) based on the normalized Shannon diversity index^21^ for each EV/RV subcomponent:

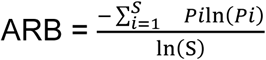

, where *Pi* represents the proportion of total EBS attributable to viral peptide *i* within a given group, and *S* is the total number of peptides in that group.

### Phage panning and ELISA

Phage panning was performed as previously described^60^. Briefly, for panning with the human scFv phage library, Maxisorp immune tubes (Thermo Scientific) were coated with a modified CE1 peptide (CSESVPALTAVETGHTS) conjugated to the C-terminus of bovine serum albumin (BSA). Three rounds of phage panning were conducted. To reduce nonspecific binding, 10% BSA was included in the blocking buffer during each round. Following panning, individual phage clones were isolated by picking single colonies and screened using a monoclonal phage Enzyme-Linked Immunosorbent Assay (ELISA). MaxiSorp 96-well plates (Fisher Scientific) were coated with antigen, and ELISA was performed according to established protocols^60^. Absorbance was measured at 450 nm using a spectrophotometer (Molecular Devices).

### Production and binding validation of the anti-CE1 scFv proteins

Production of scFvs was performed as previously described^61^. Briefly, *E. coli* HB2151 cells were transformed with phagemids encoding individual scFvs and plated on LB agar supplemented with ampicillin (100 µg/mL) and glucose (2%) at 37°C. Colonies were cultured until an optical density (OD₆₀₀) of ∼0.9. Cells were pelleted, resuspended in fresh medium without glucose, and induced with 1 mM isopropyl β-D-1-thiogalactopyranoside (IPTG) for overnight expression at 30°C. After induction, cells were collected by centrifugation and resuspended in Buffer A (1× PBS, 0.5 M NaCl, 10 mM imidazole). Periplasmic extraction of scFvs was performed using polymyxin B (Sigma-Aldrich). Cell lysates were clarified by centrifugation, and the supernatant was applied to a Ni-NTA spin column (Qiagen) for small-scale production (40 mL cultures) or to a 1 mL HisTrap column (Cytiva) for large-scale production (>500 mL cultures). Proteins were purified using an ÄKTA FPLC system (Cytiva). Eluted scFvs were dialyzed overnight against 1× PBS using 10 kDa MWCO dialysis cassettes (Thermo Fisher Scientific). Purified scFvs were subsequently evaluated by ELISA, and absorbance was measured at 450 nm with a spectrophotometer (Molecular Devices).

### Generation and binding validation of recombinant anti-CE1 antibody

The heavy- and light- chain sequence of B9 and B10 were cloned into a rabbit IgG backbone (P01870) and expressed using CHO cells to generate rB9 and rB10 (GenScript). The same variable regions were cloned to a human IgG1 backbone (P01857) and expressed using HEK293 cells to generate hB9 and hB10 (Sino Biological) all with purity >95%. Antigen specificity of these recombinant anti-CE1 antibodies were confirmed using ELISA. Briefly, a 96-well plate was first coated overnight with CE1 peptide (1 μg/mL, 100 μL per well) in carbonate-bicarbonate buffer (Sigma). After blocking with SuperBlock Buffer (Thermo Fisher), anti-CE1 antibodies and control antibody were added and incubated for two hours at room temperature. Following incubation with HRP-conjugated secondary antibody (Sigma) and TMB substrate (Thermo Fisher), antibody binding signal was measured at 450 nm using a spectrophotometer (VANTAstar, BMG LABTECH).

### HCC cell culture

Hep3B, HepG2, PLC/PRF/5, SNU182, SNU387, SNU398, SNU449 and SNU475 were obtained from ATCC, Huh1, Huh4, Huh7, and HLE were obtained from HSRRB/JCRB, SNU761 was obtained from KCLB, MHCC-97 and SMMC-7721 were obtained from Dr. Qin (Fudan University, Shanghai, China), and Hep-40 and KMCH-1 were obtained from Dr. Conner (NCI, Bethesda, USA) and Dr. Ho (NCI, Bethesda, USA), respectively.

Huh7, Huh4, PLC/PRF/5, KMCH-1, and MHCC-97 cells were maintained in Dulbecco’s modified Eagle Medium (Thermo Fisher) supplemented with 10% fetal bovine serum (FBS), 2 mM L-glutamine (Gibco), 1X penicillin and streptomycin (Gibco). Hep3B and HepG2 were cultured in Minimum Essential Medium (Thermo Fisher) supplemented with 10% FBS, 2 mM L-glutamine, 1X non-essential amino acids (Gibco), 1X sodium pyruvate (Gibco), and 1X penicillin and streptomycin. SNU387, SNU398, SNU182, SMMC-7721, HLE, and Hep-40 cells were maintained in RPMI Media (Thermo Fisher) supplemented with 10% FBS, 1X penicillin and streptomycin. SNU475, SNU449, and SNU761 were cultured in RPMI Media supplemented with 10% heat-inactivated FBS, 1X penicillin and streptomycin. All cells were maintained in a humidified 5% CO_2_ atmosphere at 37⁰C.

### Immunofluorescence staining

HCC cells were seeded on a 4-well chamber slide (Millicell) and incubated in regular culture conditions untill staining. Cells were fixed with fresh prepared 4% paraformaldehyde (Thermo) and blocked with 10% normal donkey serum (Jackson ImmunoResearch) for one hour at room temperature. rB9, rB10, and control rabbit IgG (SouthernBiotech) were diluted with PBS (Gibco) containing 2% BSA (Thermo) to a final concentration of 3 μg/mL and incubated with cells in the humidified chamber for one hour at 37 °C. After thorough wash with PBS, slides were incubated with Alexa Fluor 488-conjugated secondary antibody (1:500, Invitrogen) for 45 minutes at room temperature in the dark. Coverslips were mounted using a DAPI-containing mounting medium (Vector Labs), and images were captured using a confocal microscope (Leica Stellaris 8 FLIM) and processed using Leica LAS X software (version 1.4.5.2771).

### Surface staining and flow cytometry

HCC cells were harvested and prepared in cold FACS buffer (BD) as a single-cell suspension at a density of 1×10^6^ cells/mL. For detecting the surface binding signal of rB9, rB10 on the full panel of HCC cell lines, 18 HCC cells were incubated with rB9, rB10 and isotype control (SouthernBiotech) at 1.5 μg/mL for 30 minutes at 4 °C. For dose-response experiments, SNU475 cells were incubated with antibodies at indicated doses for 30 minutes at 4 °C. After washing, cells were incubated with Alexa Fluor 568-conjugated secondary antibody (Invitrogen) for another 30 minutes at 4 °C in the dark. Fluorescence signals were acquired on a LSRFortessa SORP cytometer (BD) and analyzed using FlowJo (version 10.10.0).

### Immunohistochemistry staining of anti-CE1 antibody in clinical HCC tissues

Formalin-fixed, paraffin-embedded tumor and adjacent non-tumor liver tissues from 68 HCC patients (TIGER-LC cohort) were deparaffinized in xylene overnight and rehydrated through a graded series of ethanol. Antigen retrieval was performed using heat-induced epitope retrieval in a citrate buffer (pH 6.2, BioGenex) for 20 minutes. Peroxidase Block (Dako) was used to quench endogenous peroxidase activity. Slides were incubated with rB9, rB10, and control antibody (SouthernBiotech) at 3 μg/mL in a humidified chamber for an hour at room temperature, followed by wash and incubation with anti-rabbit HRP Labelled Polymer (Dako) for an additional 30 minutes at room temperature. Signals were developed using DAB substrate (Dako) and counterstained with hematoxylin (Biocare Medicol). H-score was calculated by summing the products of the staining intensity and the percentage of cells at each intensity level.

### NK cell isolation and HCC co-culture

NK cells were isolated from buffy coats of healthy donors managed by the blood bank of the National Health Institute (Bethesda, MD, USA) with informed consent in accordance with the Declaration of Helsinki. The experimental protocols were reviewed and approved by the Institutional Review Board of the National Cancer Institute (Bethesda, MD, USA). Briefly, peripheral blood mononuclear cells (PBMCs) were isolated from fresh buffy coats using Ficoll (Sigma) density gradient centrifugation and cryopreserved in freezing medium (Thermo) at a density of 1×10^7^ cell/mL. NK cells were purified by negative selection using EasySep Human NK cell Isolation Kit (STEMCELL), and the isolated NK cells were cultured and expanded in CTS NK-Xpander Medium (Thermo Fisher) supplemented with IL-2 (Miltenyi).

For co-culture assays, 4,000 HCC cells per well were seeded in the 16-well E-plate (Agilent) and incubated overnight in an xCELLigence RTCA instrument. NK cells (4,000 cells per well) were added to the HCC cells together with hB9, hB10, or control human IgG (SouthernBiotech) at the indicated concentrations. HCC cell growth was monitored continuously untill the end of the experiments with the cell index calculated by the system. Control wells with only NK cells were also included, which showed no detectable signal on the E plate.

### In vivo study

Four-week-old male NUDE mice obtained from Charles River Laboratories (Rockville, MD, USA) were used in this study. Mice were group-housed (5 mice per cage) and maintained under a regular light-dark cycle altered every 12 hours with free access to water and standard mouse chow. Huh7 cells were examined regularly for Mycoplasma prior to tumor implantation. The animal study protocol (LHC-006-3) was reviewed and approved by the National Cancer Institute-Bethesda Animal Care and Use Committee (Bethesda, MD, USA).

Huh7 cells (1× 10^6^ per mouse) were injected subcutaneously in Matrigel (Corning) and DMEM (Thermo Fisher). When the tumor reached ∼100 mm^3^, mice were randomized to receive hB9, hB10 and control human IgG (SouthernBiotech) treatment (10 mg/kg). Antibody was administrated every other day (three times/week) via tail vein injection for six doses (n=9 per treatment). Tumor growth and mouse weight were then measured regularly until euthanasia. Mouse blood was obtained after euthanasia for liver function assessment. We used Alanine Transaminase Activity Assay Kit (abcam) and Aspartate Aminotransferase Activity Assay Kit (abcam) to measure the liver function of mice according to the vendor protocol. Briefly, serum was isolated from whole blood via centrafugation at 1,000g for 10 minutes at 4°C and 5 μl of serum per sample per test was used as input for both measurements. Technical duplicates were included in the test. Reagents, substrate and positive controls were prepared according to the manuals and the kinetics of the output product were measured by absorbance at OD570 nm for Alanine Transaminase Activity Assay and at 450 nm for Aspartate Aminotransferase Activity Assay (VANTAstar, BMG LABTECH). Activities of alanine transaminase and aspartate aminotransferase were computed according to the manual.

### Co-immunoprecipitation with anti-CE1 antibody

Huh7 and HLE cells were harvested and lysed in a buffer containing 150 mM NaCl (Invitrogen), 1% NP-40 (Sigma), 50 mM Tris-HCl pH=7.4 (Invitrogen), 5% (v/v) glycerol (Sigma), 1 mM EDTA (Invitrogen) supplemented with protease inhibitors (Sigma). After centrifugation at 20,000 g for 15 minutes at 4°C, lysate was collected and protein concentration was determined using BCA assay (Thermo). 500 μg of total protein was incubated with 5 μg of rB9, rB10 and control rabbit IgG overnight at 4°C with constant agitation. The immune complexes were captured using 30 μl of protein A/G magnetic beads (Thermo). Following 2-hour incubation, beads were washed three times with lysis buffer mentioned above and three times with PBS to remove non-specific bindings. A small aliquot of the complex was assessed by western blot to validate successful capture, while the remaining product was preserved in PBS and stored at −20°C for subsequent mass spectrometry analysis.

### Mass spectrometry and data analysis

Samples were solution digested with trypsin using DNA mini-prep columns, as previously described^62^. Briefly, proteins were denatured in 5% SDS, 50 mM triethylammonium bicarbonate (TEAB) pH 8.5. They were next reduced with 5 mM Tris(2-carboxyethyl)phosphine (TCEP) and alkylated with 20 mM iodoacetamide. The proteins were acidified to a final concentration of 2.5% phosphoric acid and diluted into 100 mM TEAB pH 7.55 in 90% methanol. They were loaded onto the mini-prep columns, washed four times with 100 mM TEAB pH 7.55 in 90% methanol, and digested with trypsin overnight at 37 °C. Peptides were eluted from the mini-prep columns using 50 mM TEAB, pH 8.5; 0.2% formic acid in water; and 50% acetonitrile in water. These elutions were pooled and dried by lyophilization.

Resultant peptides were analyzed using data-independent acquisition on an Orbitrap Fusion Lumos (Thermo) mass spectrometer coupled to an UltiMate 3000 RSLCnano HPLC (Thermo). Briefly, peptides were separated on a 75 µm x 25 cm, 3 µm Acclaim PepMap reverse phase column (Thermo) at 300 nL/min using and eluted directly into the mass spectrometer. Parent full-scan mass spectra collected in the Orbitrap mass analyzer set to acquire data at 120,000 FWHM resolution, followed by isolation of 4 Da non-overlapping mass windows from 430-670 for HCD fragmentation and detection in the Orbitrap at 60,000 resolution. The result data were searched against a predicted spectral library built from the same Uniprot human database using DIANN 1.9^63^ as the default settings. The intensities of identified rB9- and rB10-binders were compared with that of control antibody and a fold change >100 was used as cutoff value to define rB9- and rB10-enriched binders. The list of enriched binders was searched against a human surface protein database generated by a mass spectrometry study^27^ to identify surface proteins. Sequence details of the identified surface proteins were obtained from UniPortKB database and sequence alignment with CE1 was performed using the Align tool provided by the UniPort website.

### Anti-ASPH antibody binding ELISA

Off-the shelf anti-ASPH antibodies targeting ASPH_79-176_ (NBP2-34125) and ASPH_506-600_ (NBP2-58045) were obtained from Novus Biologicals (Centennial, CO, USA) and assessed for their recognition to CE1 peptide and recombinant ASPH antigens by ELISA. Briefly, 96 well high-bind plates (Corning) were pre-coated overnight with 100 μl of CE1 peptide or recombinant ASPH_79-176_ and ASPH_506-600_ (Novus Biologicals) at a concentration of 1 μg/mL. On the second day, plates were washed three times followed by blocking with SuperBlock Buffer. Anti-ASPH antibodies (2 μg/mL) were added and allowed to interact with the antigens for two hours at room temperature before subsequent ELISA steps. The binding signal was measured at 450 nm using a spectrophotometer (VANTAstar, BMG LABTECH).

### Recombinant ASPH antigen competition assay

Recombinant ASPH_79-176_ and ASPH_506-600_ were generated using E. coli expression system (Novus Biologicals). For CE1 binding competition assay, rB9 and rB10 at a concentration of 2 μg/mL were incubated with recombinant ASPH_79-176_ and ASPH_506-600_ at concentrations of 0, 25, 50, 100 μg/mL overnight at 4°C with continuous agitation. Meanwhile, a 96-well plate was incubated with CE1 peptide (1 μg/mL) at 4°C. On the second day, the CE1-coated plate was washed and blocked as described previously, followed by interaction with the ASPH antigen plus anti-CE1 antibody mixture for two hours at room temperature. The plate was thoroughly washed and subsequently exposed to HRP-conjugated secondary antibody and TMB substrate before being assessed by a spectrophotometer at 450 nm. For assessing the effects of ASPH antigens in B9/B10-mediated anti-HCC activities, hB9, hB10, and control IgG (5 μg/mL) were incubated with ASPH_79-176_ and ASPH_506-600_ (125 μg/mL) overnight at 4°C with continuous agitation. On the second day, these antigen-antibody mixtures were mixed with NK cells (4,000 cells per well), followed by adding to Huh7 cells pre-seeded on the E plate. The growth of Huh7 cells was continuously monitored using the xCELLigence system.

### Transcriptomic analysis of ASPH in clinical HCC samples

For comparison of the ASPH expression in tumor and non-tumor tissues, four bulk transcriptome datasets from three independent published cohorts, specifically included the LCI cohort, TCGA-LIHC cohort, and TIGER-LC cohort were used. Notably, tissues of the TIGER-LC cohorts were assessed with two different methods, specifically, 62 patients in set 1 were profiled by array and 51 patients in set 2 were profiled by RNA sequencing. Therefore, the results of the ASPH comparison were illustrated separately. For single cell study, we performed analysis using the information of ASPH expression available on the scAtlasLC. For the integrative analysis of tumor transcriptome and serological viral reactivity, we identified 97 HCC patients from TIGER cohort (set 1, n=50; set 2, n=47) with both data available. NK cell abundance was estimated using Xcell^33^, and tumors were stratified into high- and low-NK groups using a cutoff of 0.01. The expressions of ASPH and ADCC genes, including TNFRSF9, TNF, BCL2, IL21R, BIRC3, IKZF2, CCR7, CD69, MKI67, CD226, CLEC2B, CXCR4, TIA1, TNFSF10, GZMA, TLR6, TLR1, CD244, CD38, TLR3, were Z-score normalization within each dataset prior to integration. CE1 seroreactivity was defined as the mean EBS of the 40 most reactive CE1-related peptides^15^.

### Statistical Analysis

Multivariate logistic regression was performed using glm() in R. Group comparisons were conducted using two-tailed unpaired Student’s t-tests or ANOVA, as appropriate. Correlations were assessed using the Spearman rank correlation test. Survival analysis was performed using Kaplan-Meier estimates and the log-rank test. All the statistics tests were performed using GraphPad Prism Version 10.1.1 unless otherwise specified. All tests were two-sided, and *p*<0.05 was considered statistically significant.

## Data and code availability

The viral exposure profiles of the TIGER-LC cohort are available at Harvard dataverse (https://doi.org/10.7910/DVN/NIT39Z). The viral exposure profiles of the NCI-UMD cohorts are available from the corresponding author upon request. Public data sets used in this study could be downloaded from NCBI Gene Expression Omnibus or other public source under the accession number detailed in the method. All other data generated in this study are included in this published article and online supplemental materials. This paper does not report original code or generate new unique reagents.

## ACKNOWLEDGEMENTS

We are also grateful to all study participants, as well as clinicians, nurses, and study coordinators from the NCI-UMD liver study group, the TIGER-LC consortium and the NIH blood bank who helped with patient and donor enrollment. This work was supported in part by grants (ZIA-BC010313, ZIA-BC010876, ZIA-BC010877, and ZIA-BC011870 to XWW; and ZIA-BC010891 and ZIC-BC011891 to MH) from the Intramural Research Program of the Center for Cancer Research of the National Cancer Institute. The contributions of the NIH author(s) were made as part of their official duties as NIH federal employees, are in compliance with agency policy requirements, and are considered Works of the United States Government. However, the findings and conclusions presented in this paper are those of the author(s) and do not necessarily reflect the views of the NIH or the US Department of Health and Human Services.

## AUTHOR CONTRIBUTIONS

Conceptualization: MHH, XWW

Methodology: MHH, MH, XWW

Investigation: MHH, QL, LW, MF, AL, LMJ, TKM, JB, MH, JC, SR, XWW

Visualization: MHH, XWW

Funding acquisition: XWW, MH

Project administration: XWW

Supervision: XWW

Writing – original draft: MHH, XWW

Writing – review & editing: all authors

