## Extended Data Figure 1-7 and Table 1 for "Enteroviral epitope mimicry enables NK cell-mediated targeting of ASPH in hepatocellular carcinoma"

This material file includes:

Extended Data Figure 1 to 7

Extended Data Table 1

Extended Data Figure 1

A

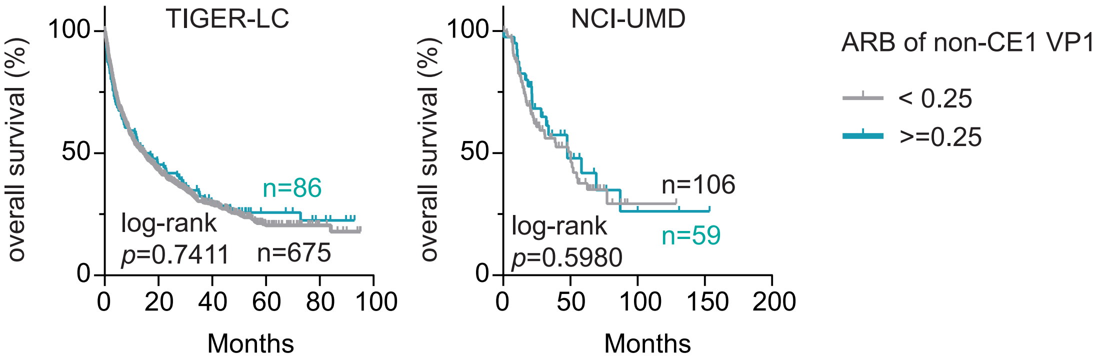


B

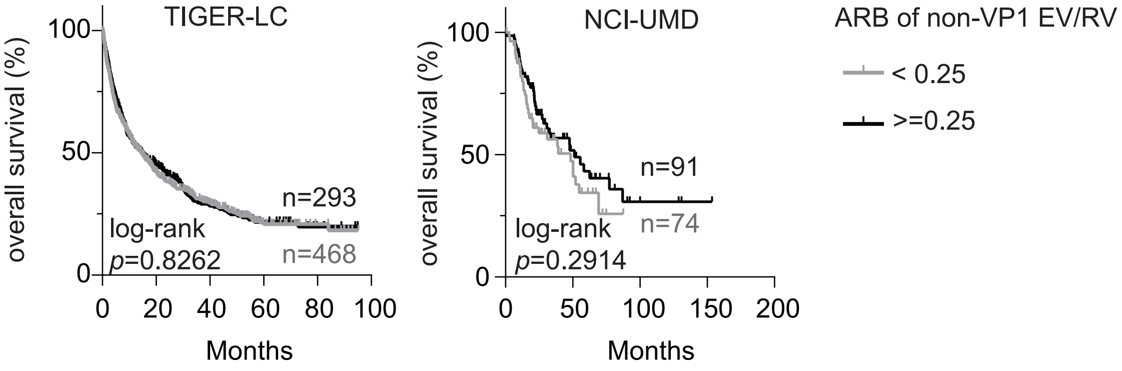

Extended Data Figure 2

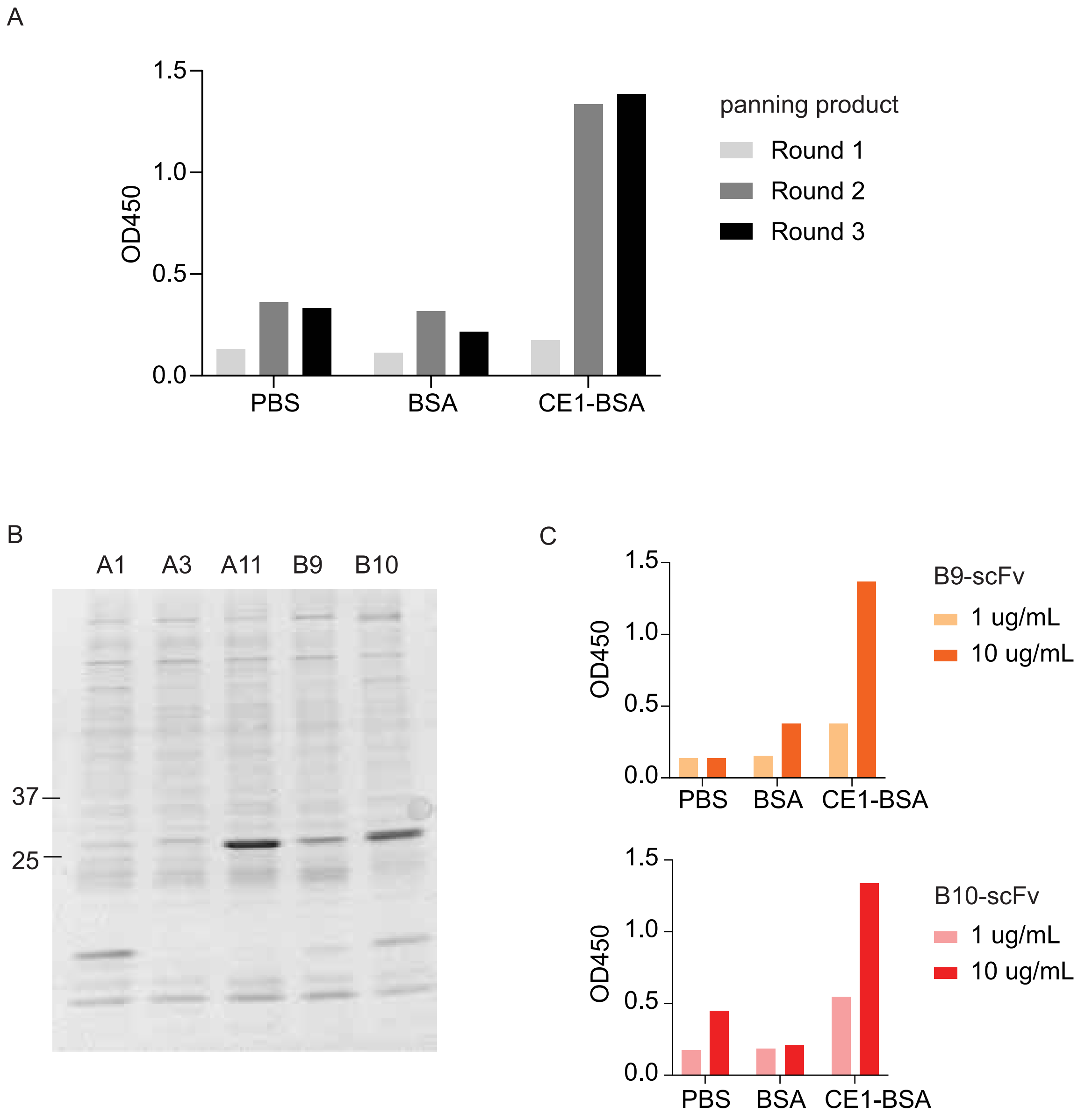


**Extended Data Figure 2. Identification of CE1-reactiving scFv from phage-display human scFv library search.**

(A)The binding between BSA-CE1-conjugated peptide and the product of the 1st, 2nd, and 3rd round of panning was measured using ELISA.

(B)Western blots of the five scFv candidates after recombinant expression.

(C)Dose-dependent CE1-binding of B9-scFv and B10-scFv.

Extended Data Figure 3

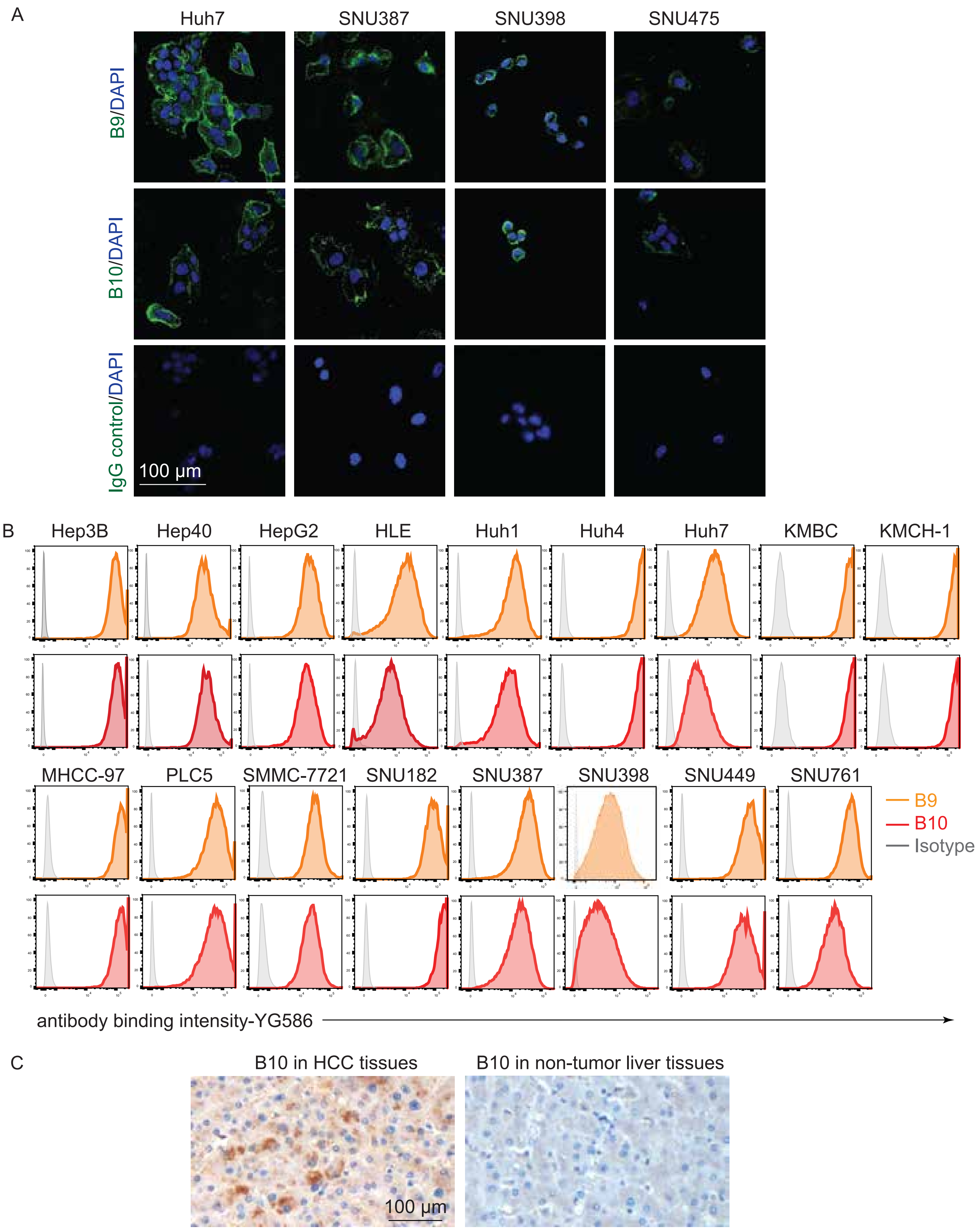

**Extended Data Figure 3. Binding of B9 and B10 to human HCC cells and clinical HCC tissues.**

(A) Representative images of immunofluorescence staining of B9, B10 and control antibody in Huh7, SNU387, SNU398, and SNU475 cells.

(B) Surface binding of B9, B10 and control antibody (2 μg/ml) to 17 HCC cell lines.

(C) Representative images of immunohistochemistry staining of B10 in clinical HCC tumor tissue and associated non-tumor tissue.

Extended Data Figure 4

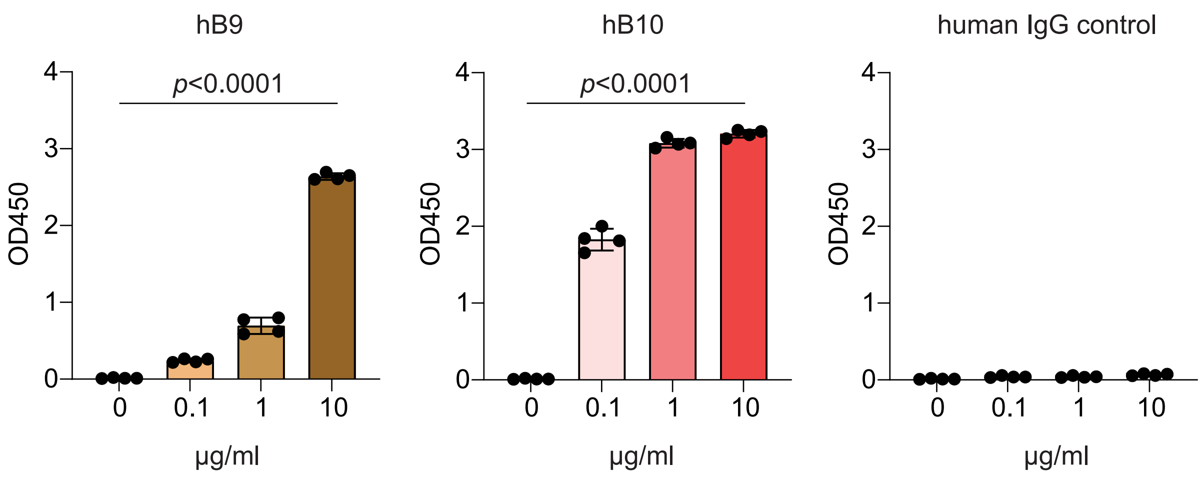


**Extended Data Figure 4. Dose-dependent CE1-binding of hB9 and hB10.**
Fully humanized B9- (hB9) and B10- (hB10) antibody and human IgG were incubated with CE1 at indicated doses for two hours and assessed by ELISA. Individual data points are plotted over mean bars with error bars representing ± standard deviation (SD). Statistical significance was determined using a one-way ANOVA test.

Extended Data Figure 5

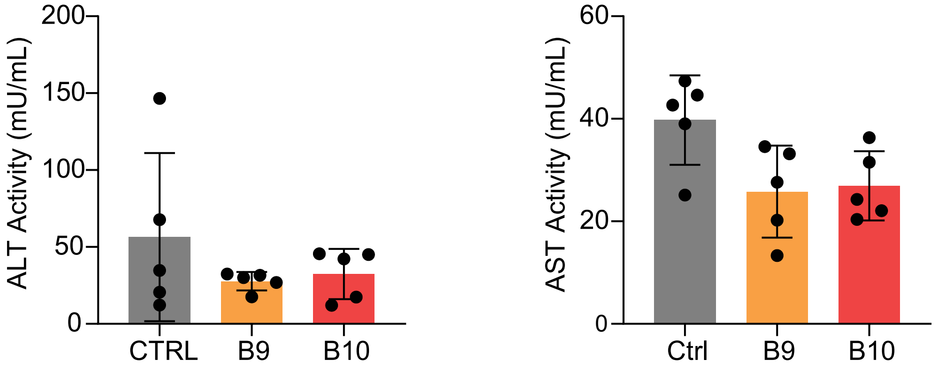


**Extended Data Figure 5. Liver function of mice receiving B9, B10 and control antibody treatment.**

Mice serums were collected upon completion of study for alanine aminotransferase and aspartate aminotransferase tests. (n=5) Individual data points are plotted over mean bars with error bars representing ± standard deviation (SD).

Extended Data Figure 6

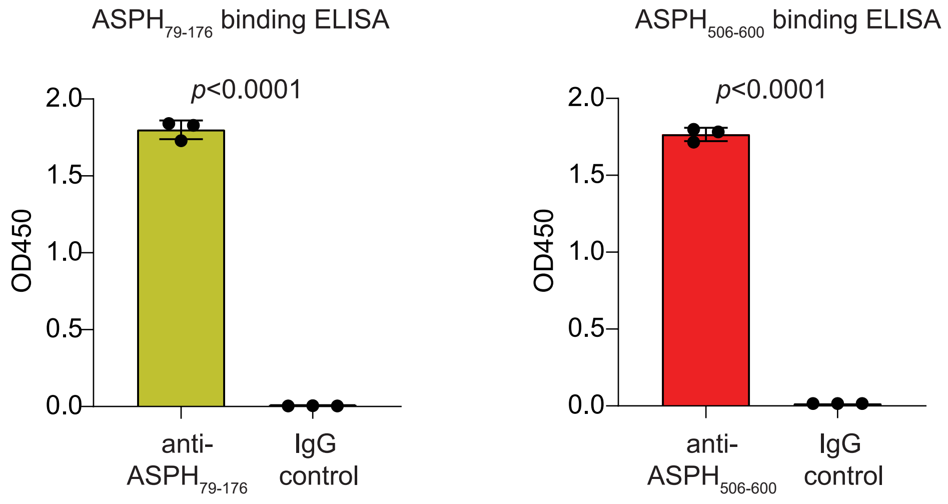


**Extended Data Figure 6. Validation of the antigenicity of recombinant ASPH antigens.** Recombinant ASPH79-176 and ASPH506-600 were incubated with the associated antibodies, namely anti- ASPH79-176 and anti-ASPH506-600, respectively, for two hours and assessed using ELISA. An isotype control was included for this experiment. (n=3) Individual data points are plotted over mean bar with error bars representing ± standard deviation (SD). Statistical significance was determined using a two-sided independent t-test.

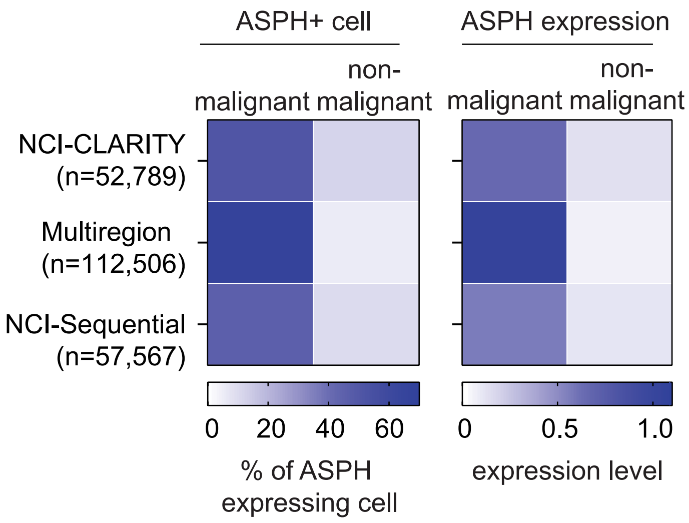


**Extended Data Figure 7. Expression of ASPH in single cell transcriptomes associated with liver cancer cells and non-malignant cells.**

Heatmap summarizes the presence of detectable ASPH and the level of ASPH expression in three independent single cell transcriptomic studies that include 31,780 malignant liver cancer cells and 157,820 non-malignant cells obtained from 101 patients.

Extended Data Table 1. Multivariate logistic regression analysis of viral serological activity and HCC diagnosis in TIGER-LC cohort.

| Viral peptide | OR (95% CI) | P value |
|---|---|---|
| CE1-VP1 | 0.90 (0.8334-0.9730) | 0.015381 |
| Non-CE1 VP1 | 0.1528 (0.0141-1.6352) | 0.121220 |
| Non-VP1 EV/RV | 0.5300 (0.0728-3.7813) | 0.527815 |
| Influenza | 5.7638 (2.0754- 16.5734) | 0.000928 |
| Corona Virus | 0.0020 (4.481486e-09 -314.2491) | 0.323984 |

EV, enterovirus; RV, rhinovirus; OR, odds ratio; CI, confidence interval
